# Cumulus cells mediate the protective effect of the marine carotenoid fucoxanthin against postovulatory oocyte aging via NRF2 signaling

**DOI:** 10.64898/2026.11.01.763385

**Authors:** Chansik Oh, Soon-Tae Kwon, Hyuk Nam Kwon, Jung-Hee Woo, Jeong Eun Lee, Sangyong Jung

## Abstract

**Background:** Oocytes awaiting insemination undergo postovulatory aging, an oxidative stress-driven loss of developmental competence. Fucoxanthin (FX), the major carotenoid of the brown alga Undaria pinnatifida, acts through NRF2, which requires a nucleus and transcription; the metaphase-II oocyte has neither, so its site of action is unresolved.

**Purpose:** To determine whether FX delays postovulatory aging of mouse oocytes and identify the mediating compartment of the cumulus–oocyte complex.

**Study design:** Ex vivo study of intact cumulus–oocyte complexes from superovulated ICR mice aged for 12 h, or 24 h for spindle analysis. Astaxanthin (AX) served as benchmark and the NRF2 inhibitor ML385 tested pathway dependence.

**Methods:** Complexes were held in M16 medium with FX (1–10 µM) or AX (5 µM). Mitochondrial membrane potential, reactive oxygen species (ROS) and glutathione were measured with JC-1, DCFH-DA and monochlorobimane; spindle organization and cumulus-cell NRF2 localization by immunofluorescence.

**Results:** At the optimal 5 µM, FX raised the JC-1 aggregate-to-monomer ratio eight-fold over untreated aged oocytes (1.24 ± 0.10 versus 0.15 ± 0.02; p < 0.05), lowered ROS to 0.18 ± 0.04-fold and raised glutathione to 1.80 ± 0.02-fold of control (both p < 0.01), comparably to AX. ML385 abolished all three effects concentration-dependently. FX increased the cumulus-cell NRF2 nuclear-to-cytoplasmic ratio (p < 0.05) and, after 24 h, nearly halved abnormal spindles (p = 0.059).

**Conclusion:** FX preserves mitochondrial and redox status in postovulatory-aged oocytes through NRF2, which translocates in cumulus cells rather than the oocyte, identifying the cumulus cell as a target for extending oocyte usability in assisted reproduction.

## 1. Introduction

Assisted reproductive technology (ART) accounts for a growing proportion of births worldwide, and because each retrieval cycle is costly, invasive and yields a limited number of oocytes, any measure that raises the usable yield of a single cycle has direct clinical value (Agarwal et al., 2022). ART implicitly assumes that retrieved oocytes are fertilized while they retain optimal developmental competence, yet an interval always separates retrieval from insemination. This interval is intrinsic to the procedure rather than a correctable lapse, arising from workflow scheduling, sperm preparation, case batching and, in donation and inter-centre programs, gamete transport. Throughout it, metaphase II (MII) oocytes remain meiotically arrested, since release from this arrest requires the calcium signal delivered at fertilization, and they are held outside the follicle and therefore without its antioxidant support. Ovulated oocytes then undergo postovulatory aging (POA), a progressive decline accompanied by rising oxidative stress, spindle disruption, chromosome misalignment, mitochondrial dysfunction, zona pellucida hardening and aberrant cortical granule dynamics, which together reduce fertilization competence and impair preimplantation development (Lord and Aitken, 2013; Miao et al., 2009). Extending the period for which retrieved oocytes can be safely held is therefore a clinically important aim.

Oxidative stress is the principal trigger of this decline. Removed from the follicular microenvironment and its antioxidant support, oocytes accumulate reactive oxygen species (ROS) while their endogenous antioxidant capacity falls, producing mitochondrial depolarization and oxidative damage to lipids, proteins and DNA (Goud et al., 2008; Miao et al., 2009). The ovulated oocyte does not face this alone. It remains enclosed by the cumulus oophorus, whose cells maintain oocyte redox homeostasis through transzonal projections and gap junctions, principally by supplying cysteine and other substrates for glutathione (GSH) synthesis that the oocyte cannot generate in sufficient amounts by itself (Maedomari et al., 2007; Zhou et al., 2010), and maintaining this coupling raises intra-oocyte GSH and lowers hydrogen peroxide (Li et al., 2016). The cumulus compartment is not a passive bystander, since cumulus cells themselves accumulate oxidative damage and undergo apoptosis as aging proceeds (Miao et al., 2009), so their supportive capacity is itself a variable that can in principle be reinforced. Consistently, delaying cumulus removal after retrieval increases the proportion of good-quality blastocysts in human ART (Mizuno et al., 2019; Patrat et al., 2012), although the optimal timing of denudation remains unestablished (Agarwal et al., 2022) and these clinical observations have not been given a mechanistic account.

Because oocytes must in any case be held in culture between retrieval and insemination, most attempts to mitigate POA have taken the holding medium as the point of intervention and relied on direct antioxidant supplementation. Melatonin and resveratrol were adopted because both are established broad-spectrum radical scavengers with documented activity in reproductive tissues, and both attenuate selected features of postovulatory aging in the mouse (Dai et al., 2017; Liang et al., 2018). Neither is in routine clinical use, and their effects are usually attributed to nonspecific modulation of the extracellular oxidative load, which leaves the endogenous antioxidant pathways of the cumulus–oocyte complex comparatively unexplored. Marine carotenoids have recently emerged as candidates here. Astaxanthin (AX) protects porcine oocytes from postovulatory deterioration and improves blastocyst yield (Jia et al., 2020), and in mice it delays POA without altering cumulus cell apoptosis or TNF-α release, an effect attributed to binding of TNFR2 on the oocyte (Wen et al., 2025). AX therefore serves both as an efficacy benchmark and, because its site of action has been localized, as a mechanistic comparator against which the target compartment of another carotenoid can be judged.

Fucoxanthin (FX) is the major carotenoid of brown macroalgae such as *Undaria pinnatifida*, which is consumed as a food in East Asia and produced at scale, so FX is available in defined, food-grade quality and has a long record of dietary exposure — practical considerations behind much of the recent interest in marine carotenoids. More importantly, FX is mechanistically distinct from both the nonspecific scavengers used previously and from AX. Rather than quenching radicals stoichiometrically, FX activates nuclear factor erythroid 2-related factor 2 (NRF2) signaling, in part through demethylation of the *Nrf2* promoter (Yang et al., 2018), and FX and its bioavailable metabolite fucoxanthinol promote NRF2 nuclear translocation and induce phase II enzymes at low micromolar concentrations (Zheng et al., 2014). Consistent with this, FX upregulates the GSH biosynthetic enzymes GCLC and GSS and thereby raises intracellular GSH through NRF2-dependent gene activation (Zheng et al., 2014) — the same metabolite that cumulus cells export to the oocyte. FX also preserves mitochondrial integrity in human ovarian granulosa cells under toxicant challenge (Elmorsy et al., 2025), indicating that follicular somatic cells respond to it. Whether FX attenuates POA, and whether such protection is cumulus-dependent, remains unknown.

We therefore hypothesized that FX delays POA by acting on NRF2-dependent antioxidant signaling in cumulus cells rather than directly on the transcriptionally quiescent oocyte. Using a mouse model of 12 h POA, we established an effective concentration of FX, compared it with AX, tested the requirement for NRF2 by pharmacological inhibition in intact complexes, quantified NRF2 nuclear translocation in cumulus cells, and assessed spindle integrity after prolonged aging. If the protective program can run only in the somatic compartment, the cumulus cell rather than the oocyte is the operative pharmacological target during postovulatory holding, and the value of any such compound would depend on keeping the cumulus investment intact — a prediction that runs counter to the practice of early denudation and is directly testable in ART.

## 2. Materials and methods

### 2.1. Chemicals and reagents

Fucoxanthin (cat. no. HY-N2302, CAS 3351-86-8, batch no. 1268697, purity 99.54% by HPLC), astaxanthin (HY-B2163), melatonin (HY-B0075) and the NRF2 inhibitor ML385 (HY-100523) were from MedChemExpress (Monmouth Junction, NJ, USA). A single batch of fucoxanthin was used throughout; the supplier’s certificate of analysis for that batch is provided in Supplementary Fig. S1. Compounds were dissolved in DMSO at 1 mM, or 10 mM, so that DMSO could be adjusted to a final 0.1% (v/v) in every group including vehicle controls. Other reagents are listed in Supplementary Methods.

### 2.2. Animals and ethics statement

Three-week-old female ICR mice were purchased from Raon Bio (Yongin, Republic of Korea) and maintained in the Laboratory Animal Research Center of CHA University under specific pathogen-free conditions at 22 ± 2 °C and 50 ± 10% relative humidity on a 12 h light/dark cycle (lights on 07:00), with sterilized food and water ad libitum. Mice were allowed to acclimatize for 24 h after arrival before any procedure and were euthanized by carbon dioxide inhalation. All experiments involving animals were conducted with the approval of the Institutional Animal Care and Use Committee of CHA University (Approval No. IACUC260114), in accordance with the Korean Animal Protection Act, and are reported in accordance with the ARRIVE 2.0 guidelines.

### 2.3. Superovulation and collection of cumulus–oocyte complexes

Mice were superovulated with 5 IU PMSG intraperitoneally at ZT11 and 5 IU hCG 48 h later. At 14 h post-hCG, cumulus–oocyte complexes (COCs) were released from the oviductal ampulla into M2 medium and retrieved mechanically, without enzymatic treatment, so that the cumulus investment remained intact. Intactness was verified under a stereomicroscope immediately after retrieval: only complexes in which the oocyte was fully enclosed by a compact, multilayered cumulus mass were used, and partially stripped, cell-shedding or damaged complexes were discarded before allocation. All treatments were applied to intact COCs throughout aging. For probe analyses and spindle immunofluorescence, oocytes were denuded identically in all groups immediately before analysis with 0.1% (w/v) hyaluronidase, and only those with an extruded first polar body, homogeneous cytoplasm and an intact plasma membrane were used as MII oocytes.

### 2.4. Postovulatory aging model and experimental design

COCs collected at 14 h post-hCG and processed without aging culture served as the non-aged reference group (Fresh). For POA, COCs were cultured in equilibrated M16 droplets under mineral oil at 37 °C under 5% CO for 12 h, or 24 h for spindle analysis. M16 is a bicarbonate-buffered embryo culture medium routinely used for mouse gamete culture, and serves here as the holding medium in which aging proceeds. All compounds were added at the onset of aging. Three designs were used. (i) *Dose-ranging and efficacy comparison*: FX at 1, 5 or 10 µM, bracketing the low micromolar concentrations at which FX promotes NRF2 nuclear translocation and phase II enzyme induction in mammalian cells (Zheng et al., 2014) while remaining below concentrations at which carotenoids become pro-oxidant, with melatonin (10 µM) as a positive control at a previously protective concentration (Dai et al., 2017); 5 µM FX was optimal (Fig. 1C, D) and used thereafter, and AX was applied at 5 µM, equimolar with FX and close to the concentration reported previously (Wen et al., 2025). (ii) *Pathway inhibition*: intact COCs co-treated with FX (5 µM) and ML385 at 0.5 or 5 µM, spanning the range over which ML385 selectively inhibits NRF2 (Singh et al., 2016). (iii) *Cumulus cell-intrinsic response*: COCs aged with vehicle, FX (5 µM) or, for transcript analysis, AX (5 µM), with a Fresh group; cumulus cells were then dissociated and processed for immunofluorescence (Section 2.6) or RNA extraction (Section 2.8). Each experiment was independently repeated three times, with COCs pooled from 5–15 mice per replicate and randomly allocated among groups within each replicate.

**Figure 1.**
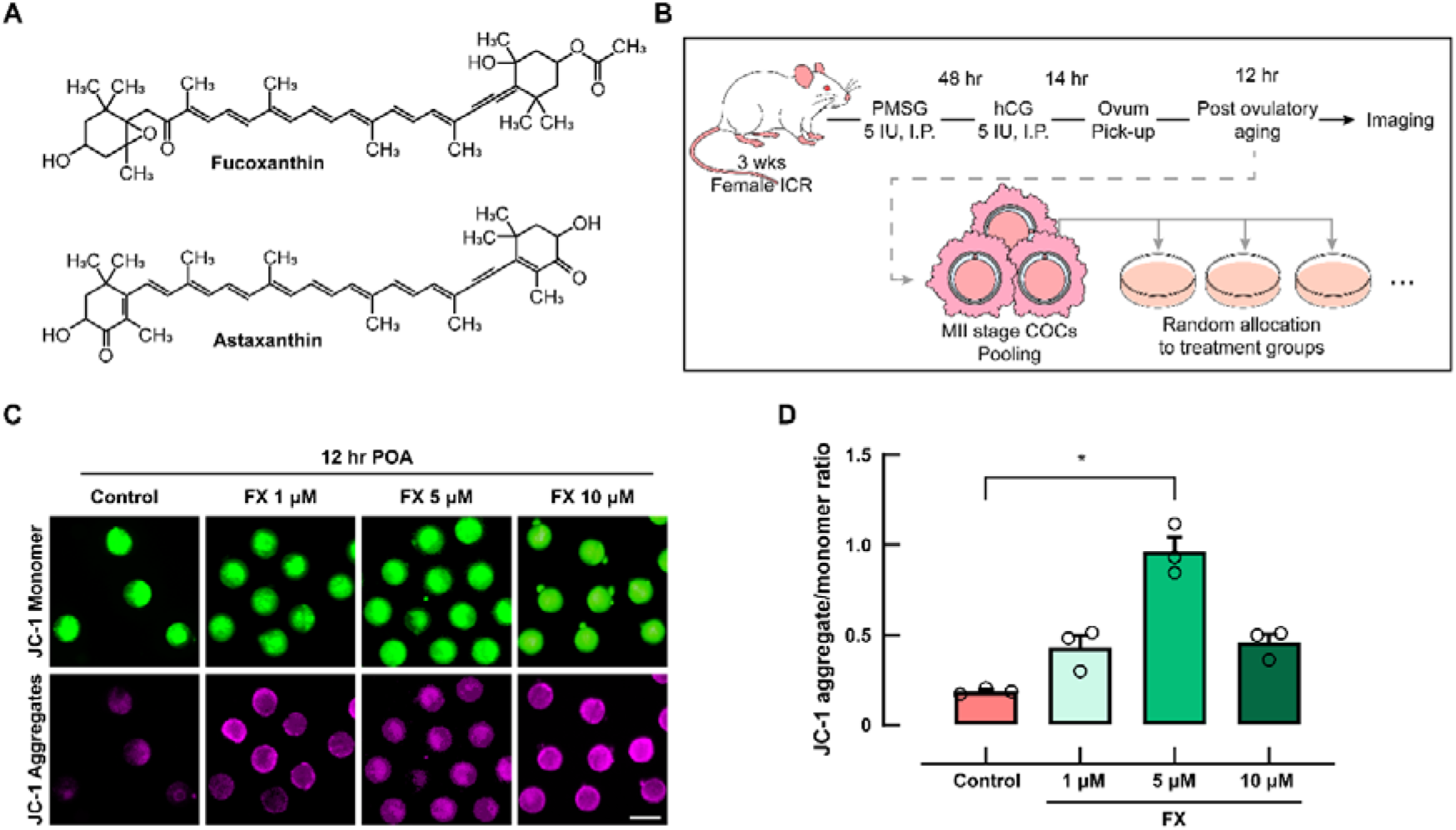
Fucoxanthin restores mitochondrial membrane potential in postovulatory-aged mouse oocytes and defines the optimal working concentration. (A) Structures of FX and AX. (B) Experimental design. Cumulus–oocyte complexes (COCs) were collected 14 h after hCG from superovulated ICR mice (5–15 mice per experiment), pooled, randomly allocated, and aged for 12 h in M16 medium; oocytes were denuded immediately before imaging. (C) JC-1 staining after 12 h of postovulatory aging (POA). Green, monomers (depolarized mitochondria); magenta, aggregates (polarized mitochondria). Control received vehicle only; FX (1, 5 or 10 µM) was added at the onset of aging. Melatonin (10 µM), included as a positive control in the same experiments, is shown in Supplementary Fig. S2. Scale bar, 100 µm. (D) Mitochondrial membrane potential (ΔΨm), expressed as the JC-1 aggregate-to-monomer ratio and quantified as described in Section 2.7. Data are mean ± SEM of three independent replicates (n = 3; 10–50 oocytes per group per replicate), and each circle represents one independent replicate. Repeated-measures one-way ANOVA with Tukey’s multiple comparisons test (*\*p <* 0.05 versus Control). The final DMSO concentration was 0.1% (v/v) in all groups.

### 2.5. Fluorescent probe analyses

Denuded oocytes were labeled for 30 min at 37 °C in the dark with DCFH-DA (10 µM) for reactive oxygen species, monochlorobimane (10 µM) for glutathione, or JC-1 (2 µg/mL) for mitochondrial membrane potential. DCFH-DA– and JC-1-labeled oocytes were imaged live, whereas monochlorobimane-labeled oocytes were fixed in 4% (w/v) paraformaldehyde for 15 min before imaging. Detection channels and full staining conditions are given in Supplementary Methods.

### 2.6. Immunofluorescence staining

Oocytes aged for 24 h, together with Fresh oocytes fixed immediately after collection, were fixed, permeabilized, blocked and immunostained for α-tubulin with DAPI counterstaining. For the cumulus cell analysis, COCs aged for 12 h were dissociated with 0.1% (w/v) hyaluronidase as for the preparation of denuded oocytes, and the released cells were attached to adhesive MAS-coated slides (HMA-MAS-11; Matsunami Glass Ind., Kishiwada, Osaka, Japan), fixed immediately and immunostained for NRF2 by the same procedure. Antibodies and dilutions are listed in Table 1; the full protocol is in Supplementary Methods.

**Table 1.**
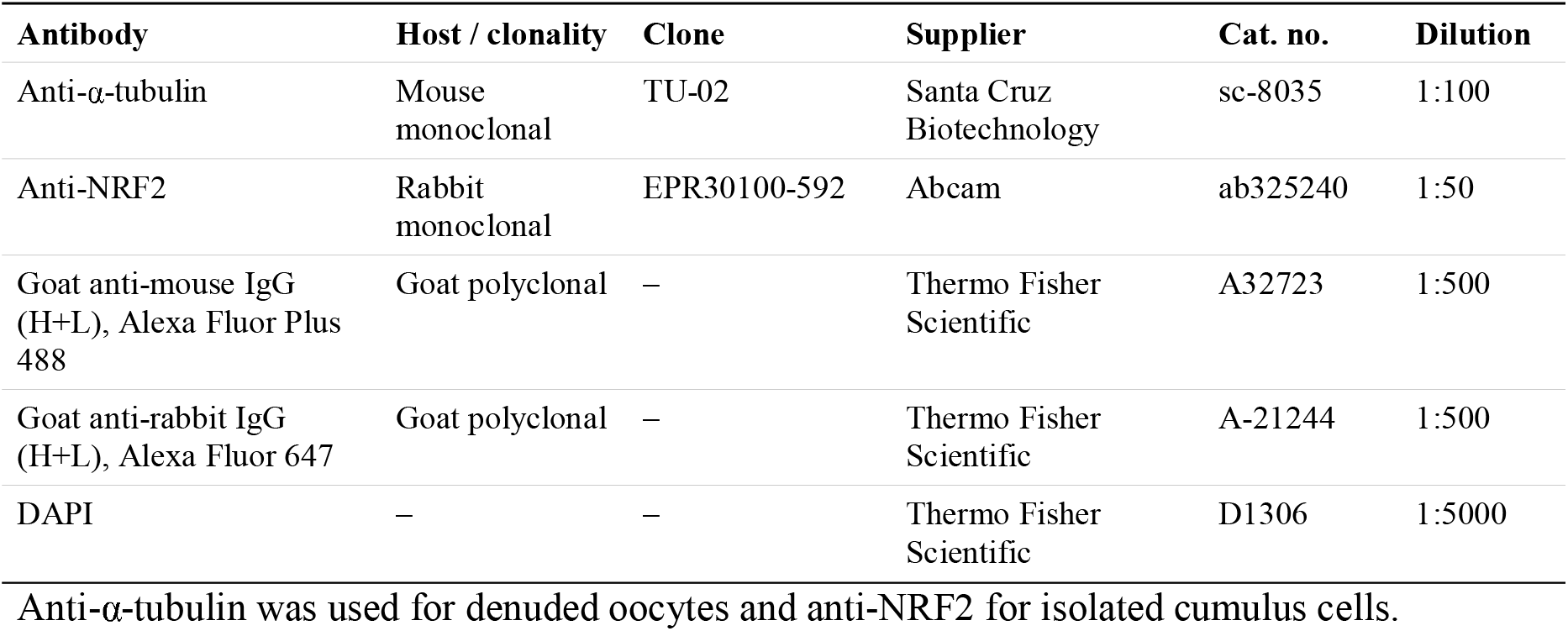
Antibodies used in this study.

### 2.7. Image acquisition and quantification

Probe signals were acquired on a RED fluorescence microscope (YourLab, Seongnam, Republic of Korea) with a ×20 objective and immunofluorescence on a BC43 confocal microscope (Andor Oxford Instruments, Belfast, Northern Ireland) with a ×60 oil-immersion objective. Within each experiment, all groups were stained and imaged in a single session on a single instrument with identical excitation intensity, exposure or gain, and detector settings; the two systems were never compared with one another, and every comparison reported here is made only among groups acquired on the same instrument in the same session. Fluorescence was quantified in Fiji/ImageJ 1.54t (National Institutes of Health, Bethesda, MD, USA). For each oocyte or cell, area, mean gray value and integrated density were obtained with the Analyze > Measure command, together with the same measurements from an adjacent cell-free region; corrected total cell fluorescence was then computed from these outputs as integrated density − (selected area × mean background gray value), and is therefore a derived quantity rather than a native ImageJ measurement. Values from all oocytes within a replicate were averaged to a single replicate mean. Mitochondrial membrane potential was expressed as the raw JC-1 aggregate (TRITC)/monomer (FITC) ratio, whereas ROS and GSH were expressed as fold of the vehicle-treated aged control mean; this rescaling does not affect the comparisons, which were performed on replicate means. NRF2 activation was expressed as the nuclear-to-cytoplasmic fluorescence intensity ratio (F^n/c^), calculated as (mean nuclear intensity − background) ÷ (mean intensity of a 1 µm perinuclear cytoplasmic annulus − background), with nuclei segmented from the DAPI channel. Meiotic spindles were scored categorically by a blinded observer, a spindle being scored abnormal only when loss of the barrel-shaped bipolar organization — disorganized, dispersed or absent microtubules, or unfocused or supernumerary poles — occurred together with chromosome misalignment, defined as displacement of one or more chromosomes from the metaphase plate by more than the width of that plate. Segmentation details are in Supplementary Methods.

### 2.8. RNA extraction and quantitative PCR

After 12 h of POA, cumulus cells were released with 0.1% (w/v) hyaluronidase as for the preparation of denuded oocytes, collected with the medium, pelleted and lysed in TRIzol (15596018, Invitrogen, Carlsbad, CA, USA). Total RNA was extracted by the TRIzol– chloroform method with glycogen as carrier, and cDNA was synthesized with the RevertAid H Minus First Strand cDNA Synthesis Kit with DNase I (K16325, Thermo Fisher Scientific, Waltham, MA, USA), which removes genomic DNA in the same reaction. Quantitative PCR was performed in technical duplicate with GoTaq qPCR Master Mix (A6001, Promega, Madison, WI, USA) on a CFX Connect Real-Time PCR Detection System (Bio-Rad Laboratories, Hercules, CA, USA) at an annealing temperature of 60 °C. Relative expression was calculated by the 2^−ΔΔCt^ method with *H2az1* (formerly *H2afz*) as the internal control and the Fresh group as the calibrator. Primer sequences are listed in Supplementary Table S1 and full procedures in Supplementary Methods.

### 2.9. Statistical analysis

All experiments were independently repeated three times, and the independent biological replicate was the experimental unit; the number of oocytes or cells per group is stated in each figure legend. Data are mean ± SEM, and normality was assessed by the Shapiro–Wilk test. Because all groups within a replicate derived from a single pool of COCs, replicates were treated as matched blocks. Blocking removes between-replicate variation, arising from differences in mouse batch, collection day and staining session, from the error term, so that treatment effects are tested against within-replicate variation only; with three replicates this materially increases sensitivity, at the cost of restricting valid comparisons to groups present within the same replicate, which was the case throughout. Comparisons among three or more groups therefore used repeated-measures one-way ANOVA with Tukey’s test, and two-group comparisons a paired *t*-test matched by batch. Where all pairwise comparisons were of interest, results are shown as compact letter notation. Proportional data were analyzed on replicate-level proportions rather than pooled counts. Analyses used GraphPad Prism 9 (GraphPad Software, San Diego, CA, USA), with *p <* 0.05 considered significant.

## 3. Results

### 3.1. FX restores mitochondrial membrane potential in postovulatory-aged oocytes

To determine whether FX protects oocytes against POA, we established an ex vivo mouse oocyte-aging model. Mitochondrial membrane potential was chosen as the primary screening readout because depolarization is among the earliest measurable events in POA, precedes overt structural damage such as spindle disassembly, and reports on the ATP-generating capacity on which the arrested oocyte depends (Takahashi et al., 2011). Structures of FX and AX are shown in Fig. 1A. MII oocytes were subjected to 12 h of POA with FX at 1, 5 or 10 µM (Fig. 1B), selected as described in Section 2.4; melatonin (10 µM) was included as a positive control in the same experiments (Supplementary Fig. S2). Membrane potential was assessed with the ratiometric probe JC-1, in which a higher aggregate-to-monomer ratio reflects greater polarization (Fig. 1C). Untreated oocytes showed a low ratio after aging, consistent with depolarization. FX raised the ratio, most markedly at 5 µM, an approximately five-fold increase over untreated aged oocytes (0.96 ± 0.08 versus 0.19 ± 0.01; \**p <* 0.05), whereas 10 µM conferred no additional benefit (Fig. 1D). Melatonin produced a robust increase, validating the responsiveness of the system to antioxidant intervention. FX therefore attenuates POA-associated mitochondrial dysfunction, and 5 µM was used thereafter.

### 3.2. FX preserves mitochondrial function and redox homeostasis in postovulatory-aged oocytes

We next compared FX with AX, which delays postovulatory aging of mouse oocytes, reducing apoptosis and spindle abnormalities and restoring embryonic development (Wen et al., 2025). Because mitochondrial and redox indices were not reported in that study, the present comparison also extends its characterization. MII oocytes were aged for 12 h with FX (5 µM) or AX (5 µM). JC-1 staining revealed pronounced depolarization in untreated aged oocytes, and both compounds raised the aggregate-to-monomer ratio significantly, FX by approximately eight-fold and AX by approximately five-fold (1.24 ± 0.10 and 0.77 ± 0.05 versus 0.15 ± 0.02; \**p <* 0.05 for each; Fig. 2A, B). Both lowered DCFH-DA fluorescence by roughly 80%, to 0.18 ± 0.04 and 0.18 ± 0.06 of control (\*\**p <* 0.01 for each; Fig. 2C, D), and raised GSH by 80% and 59%, to 1.80 ± 0.02– and 1.59 ± 0.08-fold of control (\*\**p <* 0.01 for each; Fig. 2E, F). FX and AX did not differ significantly in any parameter (*P* = 0.056, 0.973 and 0.416 for membrane potential, ROS and GSH). FX therefore preserves mitochondrial function and redox homeostasis to an extent comparable with AX. These readouts are all measured in the oocyte and cannot indicate which cell of the complex the compound acts upon, since a cumulus-mediated and an oocyte-intrinsic mechanism would produce the same oocyte phenotype; we therefore turned to the mechanism of FX action.

**Figure 2.**
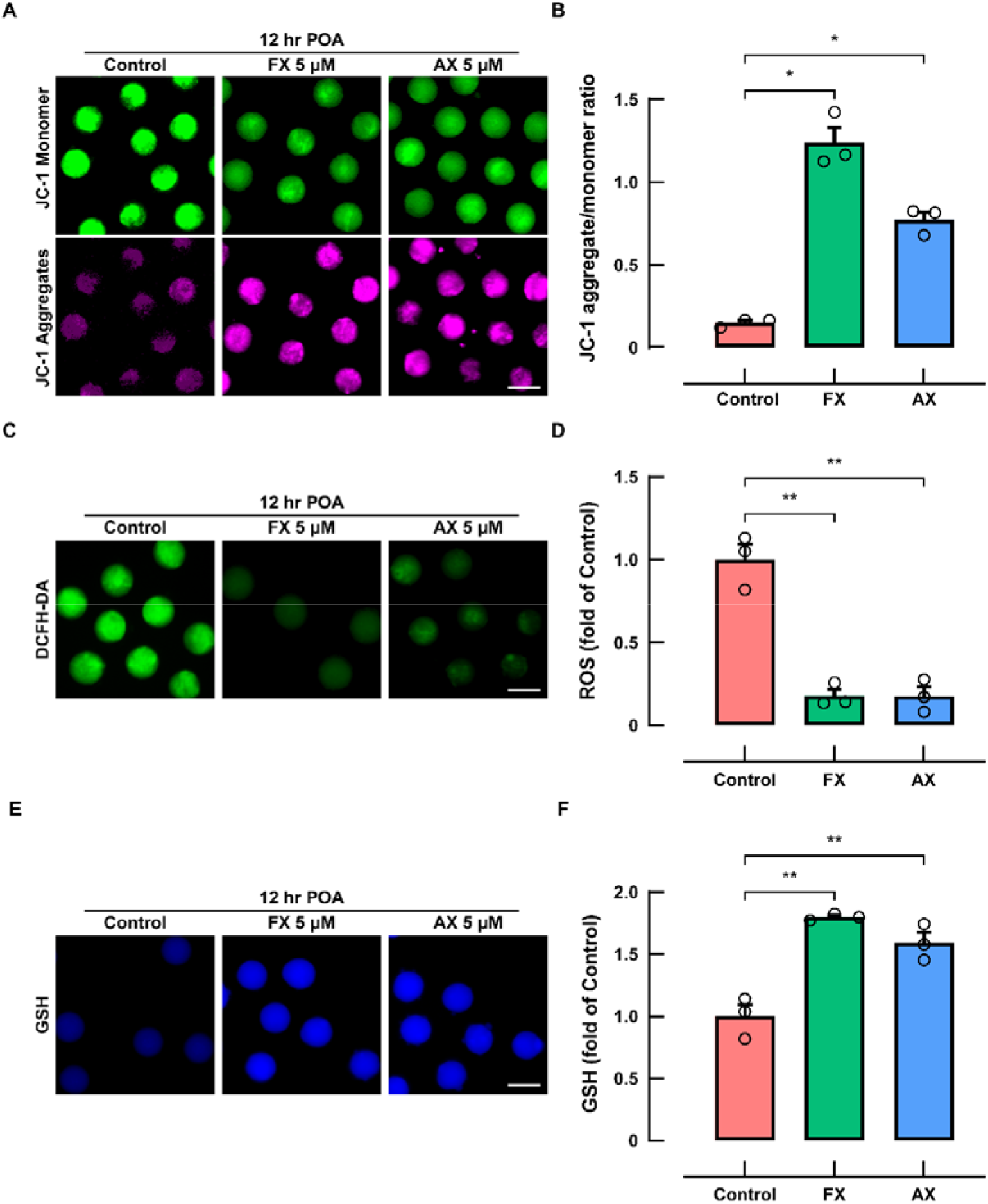
Fucoxanthin confers cytoprotection comparable to astaxanthin in postovulatory-aged mouse oocytes. (A, C, E) Representative images after 12 h of POA of JC-1 staining (A; green, monomers; magenta, aggregates), reactive oxygen species detected with DCFH-DA (C), and glutathione detected with monochlorobimane (E). FX (5 µM) or AX (5 µM) was added at the onset of aging; Control received vehicle only. Glutathione-labeled oocytes were fixed in 4% paraformaldehyde before imaging; all other panels are live. Scale bars, 100 µm. (B, D, F) Corresponding quantification, performed as described in Section 2.7; values in (D) and (F) are expressed as fold of Control. Data are mean ± SEM of three independent replicates (n = 3; 5–44 oocytes per group per replicate), and each circle represents one independent replicate. Repeated-measures one-way ANOVA with Tukey’s multiple comparisons test (\**p* < 0.05; **\*\****p <* 0.01; unmarked comparisons were not significant). DMSO was 0.1% (v/v) in all groups.

### 3.3. FX protection of postovulatory-aged oocytes requires NRF2 activity

Intact cumulus–oocyte complexes were aged for 12 h with FX (5 µM) alone or with the NRF2 inhibitor ML385 at 0.5 or 5 µM. FX raised the JC-1 ratio from 0.20 ± 0.01 to 1.81 ± 0.10 (*P* = 0.0102; Fig. 3A, B), and ML385 attenuated this concentration-dependently: at 0.5 µM the ratio fell to 0.76 ± 0.03, below the FX group (*P* = 0.0289) but still above untreated aged oocytes (*P* = 0.0006), and at 5 µM to 0.33 ± 0.04, differing from FX (*P* = 0.0194) but no longer from untreated aged oocytes (*P* = 0.122).

**Figure 3.**
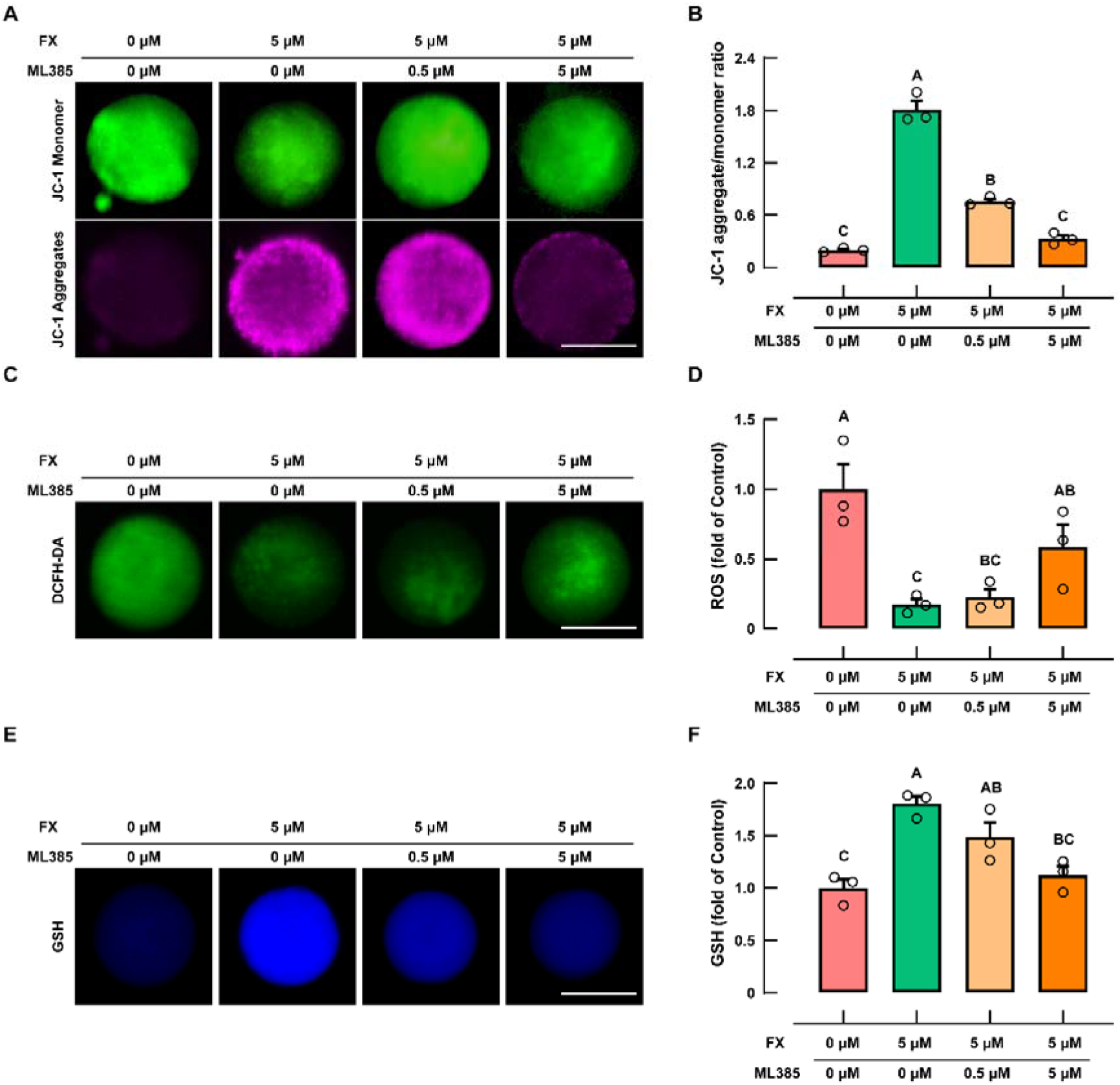
Pharmacological inhibition of NRF2 concentration-dependently abolishes the protective effect of fucoxanthin in postovulatory-aged mouse oocytes. (A, C, E) Representative images and (B, D, F) quantification of ΔΨm, reactive oxygen species and glutathione after 12 h of POA, acquired and analyzed as in Fig. 2. FX (5 µM) was given alone or together with the NRF2 inhibitor ML385 at 0.5 or 5 µM from the onset of aging; Control received vehicle only. Scale bars, 50 µm. Data are mean ± SEM of three independent replicates (n = 3; 5–54 oocytes per group per replicate), and each circle represents one independent replicate. Repeated-measures one-way ANOVA with Tukey’s multiple comparisons test for all pairwise comparisons; groups not sharing a common letter differ significantly (\**p* < 0.05).

The redox parameters showed the same dependence, with the low dose producing no statistically resolvable change and the high dose abolishing protection. FX reduced ROS to 0.17 ± 0.04 of control (*P* = 0.0042) and raised GSH to 1.81 ± 0.07-fold (*P* = 0.0029). At 0.5 µM ML385 both shifted toward the aged state but remained indistinguishable from FX alone (ROS 0.22 ± 0.06, *P* = 0.8982; GSH 1.48 ± 0.14, *P* = 0.3208), whereas at 5 µM protection was lost: ROS returned to 0.59 ± 0.16 and GSH to 1.13 ± 0.09, differing from FX (*P* = 0.0406 and 0.011) but not from untreated aged oocytes (*P* = 0.3708 and 0.7096; Fig. 3C–F). NRF2 inhibition therefore compromised, in parallel, all three protective phenotypes, indicating that intact NRF2 signaling is required and that FX acts upstream of an NRF2-dependent antioxidant program rather than as a direct radical scavenger.

### 3.4. FX promotes NRF2 nuclear translocation in cumulus cells and is associated with fewer spindle abnormalities during prolonged postovulatory aging

The requirement for NRF2 could not be met within the oocyte itself. The germinal vesicle breaks down at the resumption of meiosis, so the MII oocyte has no nuclear envelope, and transcription is globally suppressed until zygotic genome activation; there is therefore neither a nuclear compartment into which NRF2 could translocate nor transcription for it to direct. Because all treatments were applied to intact complexes, and cumulus cells remain transcriptionally active and metabolically coupled to the oocyte, we asked whether NRF2 activation occurs there. NRF2 is regulated principally through protein stability and localization rather than transcript abundance, so nuclear translocation was measured directly. After 12 h of POA, cumulus cells were dispersed, attached to adhesive slides and immunostained for NRF2. FX significantly increased the nuclear-to-cytoplasmic ratio relative to vehicle-treated aged cumulus cells (1.05 ± 0.07 versus 0.87 ± 0.04; \**p <* 0.05; Fig. 4A, B; low-magnification fields in Supplementary Fig. S3); the corresponding measurement was not attempted in the oocyte, because such a ratio is undefined in a cell without a nuclear envelope. Transcript analysis of the same cells was consistent with a mechanism operating on protein rather than message (Supplementary Fig. S4). Aging raised *Atf4* in all aged groups and *Gpx4* in the Control and FX groups relative to Fresh, and FX did not further increase either. *Nfe2l2* was not significantly raised by FX, although it was raised by AX, and the canonical targets *Hmox1* and *Nqo1* were unchanged in all groups. The only transcript significantly altered by FX was *Ptgs2*, which rose with aging and returned to the Fresh level with FX. FX therefore redistributes NRF2 protein without altering *Nfe2l2* expression; the absence of target gene induction at this single time point is considered in the Discussion.

**Figure 4.**
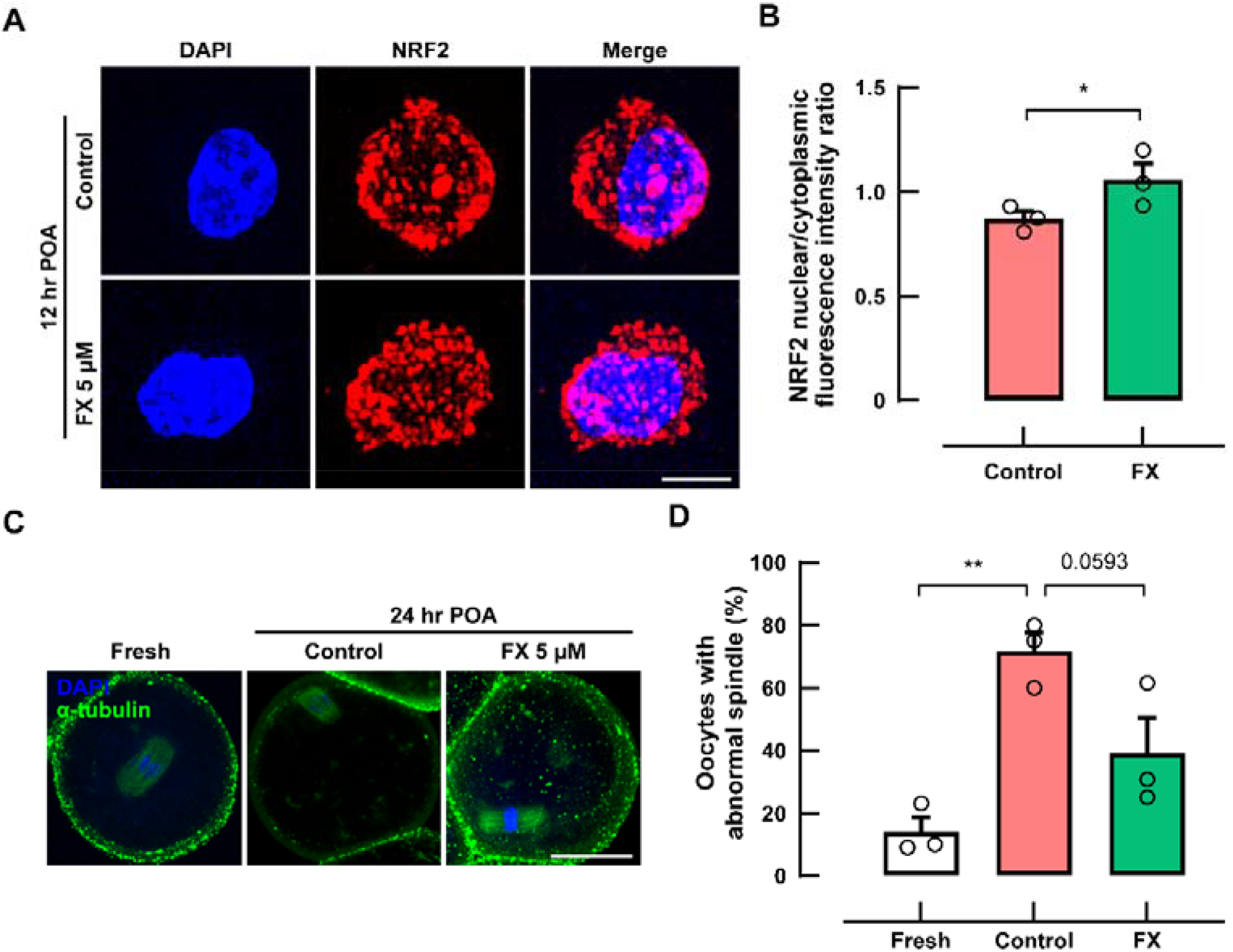
Fucoxanthin promotes NRF2 nuclear translocation in cumulus cells and is associated with fewer spindle abnormalities in postovulatory-aged oocytes. (A) NRF2 immunofluorescence (red) with DAPI counterstaining (blue) in cumulus cells dispersed from COCs after 12 h of POA. Top row, Control; bottom row, FX 5 µM. The low-magnification fields from which these cells were cropped are shown in Supplementary Fig. S3. Scale bar, 5 µm. (B) Nuclear-to-cytoplasmic fluorescence intensity ratio (F_n/c_), calculated as described in Section 2.7. Within each replicate, two fields from each of three slides prepared from separate COC pools were pooled to yield one replicate mean (5–30 cells per group per replicate), and each circle represents one independent replicate. Paired *t* test with replicates matched by batch; \**p* < 0.05. (C) Meiotic spindle after 24 h of POA: α-tubulin (green), DAPI (blue). Fresh oocytes were fixed immediately after collection. Scale bar, 50 µm. (D) Proportion of oocytes with abnormal spindles. Spindles were scored by an observer blinded to treatment using the criteria defined in Section 2.7, an abnormal spindle being one in which loss of the barrel-shaped bipolar organization occurred together with chromosome misalignment, the latter defined as displacement of one or more chromosomes from the metaphase plate by more than the width of that plate. Within each replicate, Fresh, Control and FX oocytes were derived from the same collection pool (10–13 oocytes per group per replicate; 34, 32 and 35 in total). Data are mean ± SEM of three independent replicates (n = 3), and each circle represents one independent replicate; repeated-measures one-way ANOVA with Tukey’s multiple comparisons test (\*\**p <* 0.01; the exact *P* value is shown for Control versus FX). FX (5 µM) was added at the onset of aging and DMSO was 0.1% (v/v) in all groups.

We next asked whether FX preserves meiotic spindle integrity during prolonged aging. Oocytes were aged for 24 h, when spindle deterioration is nearly complete in this model, with Fresh, aged control and FX-treated oocytes drawn from the same collection pool within each replicate. Spindles were scored against the criteria defined in Section 2.7 by an observer blinded to group allocation. Only 14.1 ± 4.5% of Fresh oocytes were abnormal, whereas aging raised this five-fold, to 71.7 ± 6.0% (*P* = 0.0049; Fig. 4C, D). FX nearly halved the proportion, to 38.8 ± 11.4%; although this approached but did not reach significance (*P* = 0.0593), FX-treated oocytes were no longer distinguishable from Fresh oocytes (*P* = 0.1396), and the direction of the effect matched the mitochondrial and redox data. Together with the inhibition experiments, these findings support a model in which FX engages NRF2-dependent signaling within the cumulus cell compartment and thereby preserves oocyte integrity during POA.

## 4. Discussion

Postovulatory aging compromises fertilization and subsequent embryonic development (Lord and Aitken, 2013; Miao et al., 2009), yet strategies capable of extending the functional lifespan of ovulated oocytes remain limited. Here we identify FX as a marine carotenoid that attenuates several hallmarks of POA and provide evidence linking its protective activity to NRF2 signaling within the cumulus cell compartment. FX preserved mitochondrial membrane potential, suppressed ROS accumulation and increased GSH in aged oocytes, to an extent comparable at the phenotypic level with AX; pharmacological inhibition of NRF2 progressively abolished these benefits; FX promoted NRF2 nuclear accumulation in cumulus cells; and FX showed a tendency to preserve spindle integrity.

After ovulation, MII oocytes remain arrested while accumulating ROS and depleting antioxidant capacity, with consequent mitochondrial depolarization and damage to proteins, lipids, DNA and the meiotic apparatus (Goud et al., 2008; Takahashi et al., 2011). Because oocytes depend on maternally inherited mitochondrial pools and cannot replace damaged organelles during meiotic arrest, ROS-driven loss of membrane potential establishes a self-amplifying cycle. Aged control oocytes showed exactly this phenotype, and FX reversed all three indices simultaneously, indicating coordinated preservation of redox and mitochondrial homeostasis rather than an effect on a single marker. Protection was maximal at 5 µM and not increased at 10 µM; since carotenoids can perturb redox balance at higher concentrations, this non-linear response is itself more consistent with regulated signaling than with stoichiometric radical quenching.

Comparison with AX supports this interpretation while raising a further question. Both compounds converged on the same oocyte phenotype, yet phenotypic similarity does not imply mechanistic equivalence. AX was recently reported to extend the functional lifespan of mouse oocytes through a mechanism attributed to TNFR2 in the oocyte, without altering cumulus cell apoptosis or TNF-α release (Wen et al., 2025). Our data suggest that FX reaches a comparable endpoint through NRF2 signaling in the surrounding somatic compartment. That two structurally related xanthophylls should converge from different points within the same complex is not paradoxical, since the complex is a functional unit with an established capacity to transfer reducing equivalents inward. FX is appreciably more polar than AX, which may favor retention in the outer cumulus layers, but we regard this as a hypothesis for direct measurement of compound distribution rather than an explanation established here. The requirement for NRF2 provides mechanistic support. NRF2 coordinates transcription of genes governing GSH synthesis, antioxidant defense and detoxification (Cuadrado et al., 2019; Yamamoto et al., 2018), and inhibition with ML385 (Singh et al., 2016) progressively attenuated all three protective phenotypes. Coordinated reversal of functionally related but independent endpoints argues against nonspecific radical scavenging, with the pronounced effect on GSH consistent with the established control of GSH biosynthetic genes by NRF2. Where this response occurs is constrained by the biology of the MII oocyte, which lacks both a nuclear envelope and, until zygotic genome activation, active transcription (Bouniol-Baly et al., 1999). Any NRF2-dependent response during aging must therefore arise in a transcriptionally competent cell, and in the ovulated complex the only such cell is the cumulus cell, which remains metabolically coupled to the oocyte through transzonal projections and gap junctions (Martinez et al., 2023). Our finding that FX increased the nuclear-to-cytoplasmic NRF2 ratio in cumulus cells provides a cellular basis for the pharmacological data.

This interpretation is particularly relevant to GSH, the metabolite whose synthesis in the oocyte depends most directly on the cumulus compartment (Maedomari et al., 2007; Zhou et al., 2010) and the parameter most strongly affected by FX here. NRF2 activation in cumulus cells could enhance cysteine availability and GSH biosynthetic capacity, as documented for NRF2-expressing glia supporting adjacent neurons (Shih et al., 2003). Our experiments do not, however, demonstrate transfer of GSH or its precursors, and two alternatives remain open: paracrine transfer through secreted factors or extracellular vesicles, and a purely local effect in which the cumulus mass lowers the oxidative load reaching the oocyte. Gap-junctional coupling also declines around ovulation, so its contribution during postovulatory holding cannot be assumed from studies of the maturing complex.

Whether these improvements preserve meiotic architecture is functionally the decisive question. Prolonged aging markedly increased the proportion of abnormal spindles, and FX lowered it. Although the difference from untreated aged controls approached but did not reach significance (*P* = 0.0593), FX-treated oocytes were no longer distinguishable from freshly collected oocytes. This relationship is mechanistically plausible: spindle assembly and maintenance in the MII oocyte depend directly on mitochondria-derived ATP, and oxidative challenge disassembles the MII spindle through a fall in membrane potential and cytoplasmic ATP rather than through a direct action on tubulin (Zhang et al., 2006), while agents that restore mitochondrial function in stressed oocytes also restore spindle organization and chromosome alignment (Guo et al., 2020). Improved mitochondrial and antioxidant status could therefore contribute to meiotic integrity, but additional replicates and direct assessment of aneuploidy will be required before this can be concluded.

Several limitations should be stated. ML385 was added to the intact complex and therefore inhibits NRF2 in both cell types; the experiment establishes that NRF2 activity is required but does not by itself assign that requirement to the cumulus cell, for which conditional deletion of *Nfe2l2* in cumulus or granulosa cells would be needed. We did not include an inhibitor-only group, although the high-dose combination returned the redox indices to the level of the vehicle-treated aged control rather than below it. Our argument concerns the canonical, transcription-dependent function of NRF2, and a transcription-independent action within the oocyte cannot be excluded. With three replicates the power to resolve small differences is limited; the FX–AX comparison in membrane potential (*P* = 0.056) and the spindle comparison (*P* = 0.0593) both fall on the threshold and are not treated as evidence of equivalence. The most substantial gap concerns the downstream half of the mechanism: FX increased nuclear NRF2 but did not induce *Hmox1* or *Nqo1*, so the chain from translocation to target gene output is not closed by our data. A single 12 h endpoint may fall outside the window in which these transcripts peak, and the between-replicate variance of *Hmox1* was large enough that a moderate induction would not have been resolved; a time course of NRF2 target transcripts and proteins, with direct measurement of GCLC, GSS and GSH in cumulus cells, is the necessary next step, as is calibration of the nuclear-to-cytoplasmic ratio against a canonical activator such as sulforaphane. Spindle integrity was assessed by blinded categorical scoring, which is conventional but less sensitive than continuous measurement; spindle dimensions, chromosome dispersion and kinetochore-based aneuploidy analysis would provide a graded readout. Finally, FX is deacetylated to fucoxanthinol by esterases in other tissues and the active species here is unknown, and the study addressed cellular indices only, so fertilization and developmental outcomes remain to be determined.

These limitations notwithstanding, the findings have implications for ART. Most antioxidant strategies have sought to protect the oocyte by modifying the extracellular redox environment (Dai et al., 2017; Liang et al., 2018); our data suggest instead reinforcing the endogenous protective capacity of the complex itself. This also offers a mechanistic account of observations so far treated as empirical. Delaying cumulus removal after retrieval raised the proportion of good-quality blastocysts (Mizuno et al., 2019), and a longer retrieval-to-denudation interval improved fertilization whereas a longer denudation-to-injection interval reduced it (Patrat et al., 2012), yet the optimal timing of denudation is still described as unestablished (Agarwal et al., 2022). If the program that limits postovulatory deterioration can run only in the somatic compartment, removing that compartment removes the capacity to run it. The corollary is testable, and a preliminary experiment is consistent with it: oocytes exposed to FX throughout 24 h of aging showed abnormal spindles in about half of cases when aged inside intact complexes, but in every oocyte examined when denuded before culture, the latter proportion matching that of vehicle-treated denuded controls (Supplementary Fig. S5). That experiment was performed once and its control arm consisted of denuded oocytes only, so it is reported without statistical analysis and as preliminary. We do not propose a change in clinical practice from a murine model, but the clinical observations and the present mechanism are mutually supporting, and whether FX can extend this window in human oocytes remains to be determined.

## 5. Conclusion

FX, a chemically defined carotenoid of brown macroalgae, attenuates postovulatory oocyte aging, preserving mitochondrial polarization, limiting ROS accumulation, enhancing GSH availability and showing a tendency to maintain spindle integrity. Its protective activity is linked to NRF2-dependent signaling in the cumulus cell compartment, which — because the MII oocyte lacks a nuclear envelope and active transcription — is the only cell of the complex able to execute such a program. These findings highlight the cumulus cell as a pharmacologically accessible regulator of oocyte resilience.

## CRediT authorship contribution statement

C. Oh: Conceptualization, Methodology, Investigation, Formal analysis, Visualization, Writing – original draft.

S.-T. Kwon: Methodology, Investigation, Resources, Writing – review and editing.

H.N. Kwon: Methodology, Investigation, Resources, Writing – review and editing.

J.-H. Woo: Methodology, Investigation, Resources, Writing – review and editing.

J.E. Lee: Conceptualization, Supervision, Project administration, Writing – review and editing.

S. Jung: Conceptualization, Supervision, Funding acquisition, Project administration, Writing – review and editing.

All authors read and approved the final manuscript.

## Ethics statement

All animal procedures were reviewed and approved by the Institutional Animal Care and Use Committee of CHA University (approval number IACUC260114) and were carried out in accordance with institutional and national guidelines for the care and use of laboratory animals.

## Data availability

Data will be made available on request. All data supporting the conclusions are presented in the article and its supplementary material; raw imaging files and the source data underlying each figure are available from the corresponding author on reasonable request.

## Declaration of generative AI and AI-assisted technologies in the writing process

During the preparation of this work the authors used Claude AI in order to improve language and readability. After using this tool, the authors reviewed and edited the content as needed and take full responsibility for the content of the publication.

## Abbreviations

ART: assisted reproductive technology
AX: astaxanthin
COC: cumulus–oocyte complex
DCFH-DA: 2′,7′-dichlorodihydrofluorescein diacetate
FX: fucoxanthin
GCLC: glutamate cysteine ligase catalytic subunit
GSH: glutathione
GSS: glutathione synthetase
hCG: human chorionic gonadotropin
MII: metaphase II
NRF2: nuclear factor erythroid 2-related factor 2
PMSG: pregnant mare serum gonadotropin
POA: postovulatory aging
ROS: reactive oxygen species
TNFR2: tumor necrosis factor receptor 2
ZT: Zeitgeber time
ΔΨm: mitochondrial membrane potential

---

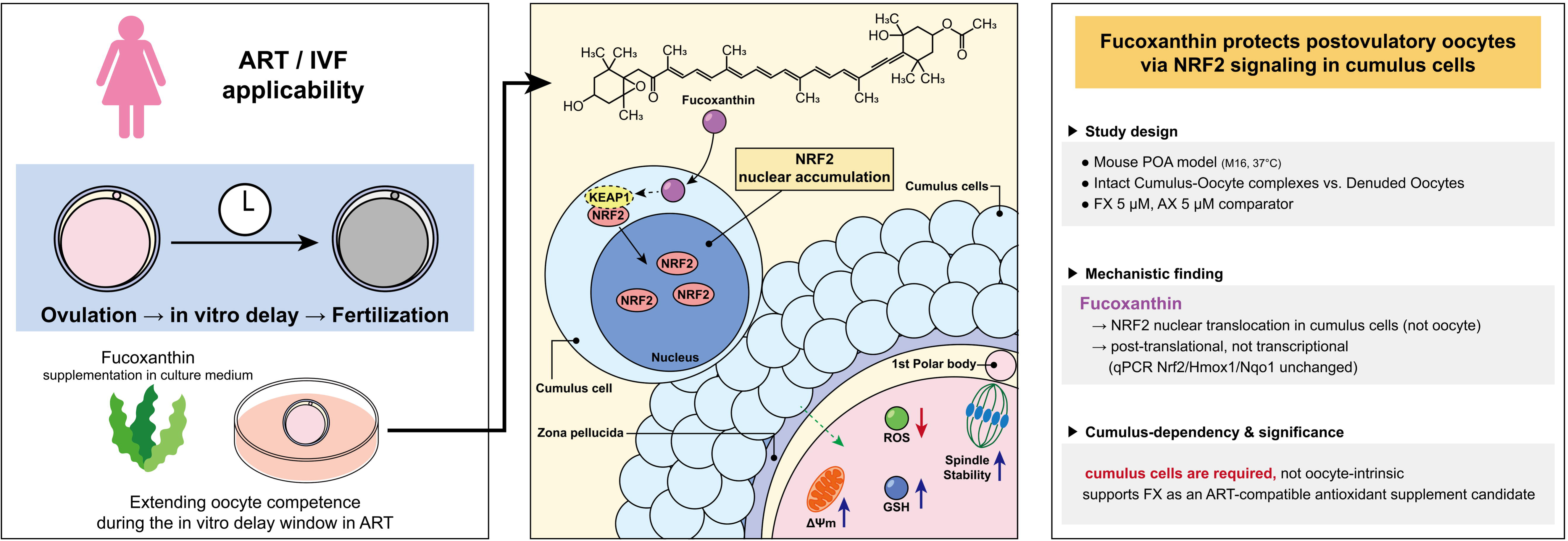

