## Supplementary material for "Cumulus cells mediate the protective effect of the marine carotenoid fucoxanthin against postovulatory oocyte aging via NRF2 signaling": Tables

Table 1. Antibodies used in this study.

| **Antibody** | **Host / clonality** | **Clone** | **Supplier** | **Cat. no.** | **Dilution** |
| --- | --- | --- | --- | --- | --- |
| Anti-α-tubulin | Mouse monoclonal | TU-02 | Santa Cruz Biotechnology | sc-8035 | 1:100 |
| Anti-NRF2 | Rabbit monoclonal | EPR30100-592 | Abcam | ab325240 | 1:50 |
| Goat anti-mouse IgG (H+L), Alexa Fluor Plus 488 | Goat polyclonal | – | Thermo Fisher Scientific | A32723 | 1:500 |
| Goat anti-rabbit IgG (H+L), Alexa Fluor 647 | Goat polyclonal | – | Thermo Fisher Scientific | A-21244 | 1:500 |
| DAPI | – | – | Thermo Fisher Scientific | D1306 | 1:5000 |

Anti-α-tubulin was used for denuded oocytes and anti-NRF2 for isolated cumulus cells.
