## Supplementary data for "Cumulus cells mediate the protective effect of the marine carotenoid fucoxanthin against postovulatory oocyte aging via NRF2 signaling"

**Supplementary Methods**

**Reagents**

Pregnant mare serum gonadotropin was obtained from Daesung Microbiological Labs (Uiwang, Republic of Korea). Human chorionic gonadotropin (C1063), M2 medium (M7167), hyaluronidase from bovine testes (H4272), monochlorobimane (69899), 2′,7′-dichlorodihydrofluorescein diacetate (287810), polyvinyl alcohol (341584) and Triton X-100 (T8787) were from Sigma-Aldrich (St. Louis, MO, USA); M16 medium (MR-016-D) and FluorSave Reagent (345789) from Merck Millipore (Burlington, MA, USA); bovine serum albumin (10735086001) from Roche (Basel, Switzerland); 4% paraformaldehyde (P2031) from Biosesang (Yongin, Republic of Korea); goat serum (BGN-17) from Dojindo Korea (Seoul, Republic of Korea); and JC-1 (65-0851-38) from eBioscience (San Diego, CA, USA). DMSO was from LPS Solution (DMSO100, Daejeon, Republic of Korea). Aliquots of all test compounds were stored at −80 °C protected from light. Unless otherwise specified, remaining reagents were from Sigma-Aldrich.

**Postovulatory aging culture**

M16 medium droplets (70 µL) were covered with mineral oil and equilibrated at 37 °C under 5% CO₂ in humidified air for 12 h before use; mineral oil was equilibrated under identical conditions before use. Denudation was performed by exposure to 0.1% (w/v) hyaluronidase in M2 medium for 1–3 min, followed immediately by three washes in fresh M2 medium and gentle pipetting with a glass capillary.

**Fluorescent probe analyses**

Oocytes were incubated with the respective probe in M2 medium at 37 °C for 30 min in the dark and washed three times in PBS containing 0.1% (w/v) polyvinyl alcohol. DCFH-DA signal was acquired in the FITC channel and monochlorobimane in the DAPI channel; JC-1 was acquired in both the FITC (monomer) and TRITC (aggregate) channels. Within each experiment, all groups were stained and imaged in the same session using identical acquisition settings.

**Immunofluorescence staining**

Oocytes were fixed for 30 min in 4% (w/v) paraformaldehyde in PBS containing 0.1% (w/v) polyvinyl alcohol, pre-warmed to 37 °C and allowed to equilibrate to room temperature. Samples were permeabilized with 0.3% (v/v) Triton X-100 for 15 min and blocked overnight at 4 °C in PBS containing 5% (v/v) goat serum, 1% (w/v) bovine serum albumin, 0.05% (v/v) Triton X-100 and 0.1% (w/v) polyvinyl alcohol, then incubated overnight at 4 °C with the primary antibody. Bound antibody was detected with the corresponding Alexa Fluor-conjugated secondary antibody for 1 h at room temperature, together with DAPI. All steps were separated by three 15 min washes in PBS–0.1% polyvinyl alcohol. Oocytes were mounted in FluorSave Reagent and examined by confocal or widefield fluorescence microscopy; the same modality was used for all groups within a replicate. Cumulus cells were washed three times in M2 medium and allowed to attach to adhesive slides for 5 min at 37 °C before immediate fixation, then processed identically.

**Image quantification**

For NRF2, nuclear regions were segmented from the DAPI channel and the perinuclear annulus was generated as a 1 µm band immediately outside the nuclear boundary, from which the nuclear masks of all cells in the field were subtracted so that adjacent nuclei did not contribute. Background was measured in cell-free regions of the same image. Within each replicate, two fields were analyzed from each of three slides prepared from separate COC pools, and all cells were pooled to yield a single replicate mean. Meiotic spindles were scored categorically by an observer blinded to group allocation, using the criteria defined in Section 2.7 of the main text. An oocyte was scored as abnormal only when loss of the typical barrel-shaped bipolar spindle — disorganized, dispersed or absent microtubules, or unfocused or supernumerary poles — occurred together with chromosome misalignment, the latter defined as displacement of one or more chromosomes from the metaphase plate by more than the width of that plate.

**RNA extraction and quantitative PCR**

Cumulus cells were collected with the medium, pelleted at 300 × g for 3 min, lysed in 500 µL TRIzol (15596018, Invitrogen, Carlsbad, CA, USA) and stored at −80 °C until extraction; all samples of a given replicate were extracted together. RNA was isolated by the standard TRIzol–chloroform method with glycogen (20 µg; R0551, Thermo Fisher Scientific) as carrier and overnight ethanol precipitation at −20 °C, washed in 75% ethanol prepared in DEPC-treated water, dissolved in 12 µL RNase-free water and quantified by NanoDrop spectrophotometry. Complementary DNA was synthesized with the RevertAid H Minus First Strand cDNA Synthesis Kit with DNase I (K16325, Thermo Fisher Scientific), and quantitative PCR was performed with GoTaq qPCR Master Mix (A6001, Promega, Madison, WI, USA), following the manufacturers' instructions in each case.

**Supplementary Table**

Table S1. Primer sequences used for quantitative PCR.

| **Gene Symbol** | **Accession no.** | **Forward Primer (5′→3′)** | **Reverse Primer (5′→3′)** | **Amplicon size (bp)** |
| --- | --- | --- | --- | --- |
| *Gpx4* | NM_008162 | GAGGAGCCCCAGGTGATAGA | AGGCTGGTTTTCAGGCAGAC | 146 |
| *Atf4* | NM_009716 | CTGCCTTCTCCAGGTGGTTC | TGGCTGCTGTCTTGTTTTGC | 139 |
| *Hspa5* | NM_022310 | TCAGGAACTTTCGTTGGGAGA | ACCCCAAGACATGTGAGCAA | 144 |
| *Nfe2l2* | NM_010902 | GGCACCATTTGGGAGAGTTT | TCTGGTTAGTTTTCAATAGTTTGCT | 127 |
| *Hmox1* | NM_010442 | GTTGACTGACCACGACTGCT | ACTCTGGTAGAACTGGGGAAA | 117 |
| *Nqo1* | NM_008706 | AGAGAGTGCTCGTAGCAGGAT | GTGGTGATAGAAAGCAAGGTCTT | 94 |
| *Ptgs2* | NM_011198 | TCAGGACTACTATGTACCTCCCC | ATATTTCTAGCATCTTGGTAGGGC | 122 |
| *H2az1* | NM_016750 | AGTGGGTTTTGATTGAGTTGACTG | ACCAGAGTGGAAACAATGTCTGT | 113 |

*H2az1* (formerly *H2afz*) was used as the internal control gene. All reactions were performed at an annealing temperature of 60 °C. Primers were designed using Primer-BLAST (NCBI) against the indicated reference sequences and span exon–exon junctions. Accession numbers refer to NCBI RefSeq entries for *Mus musculus*.

**Supplementary Figures**

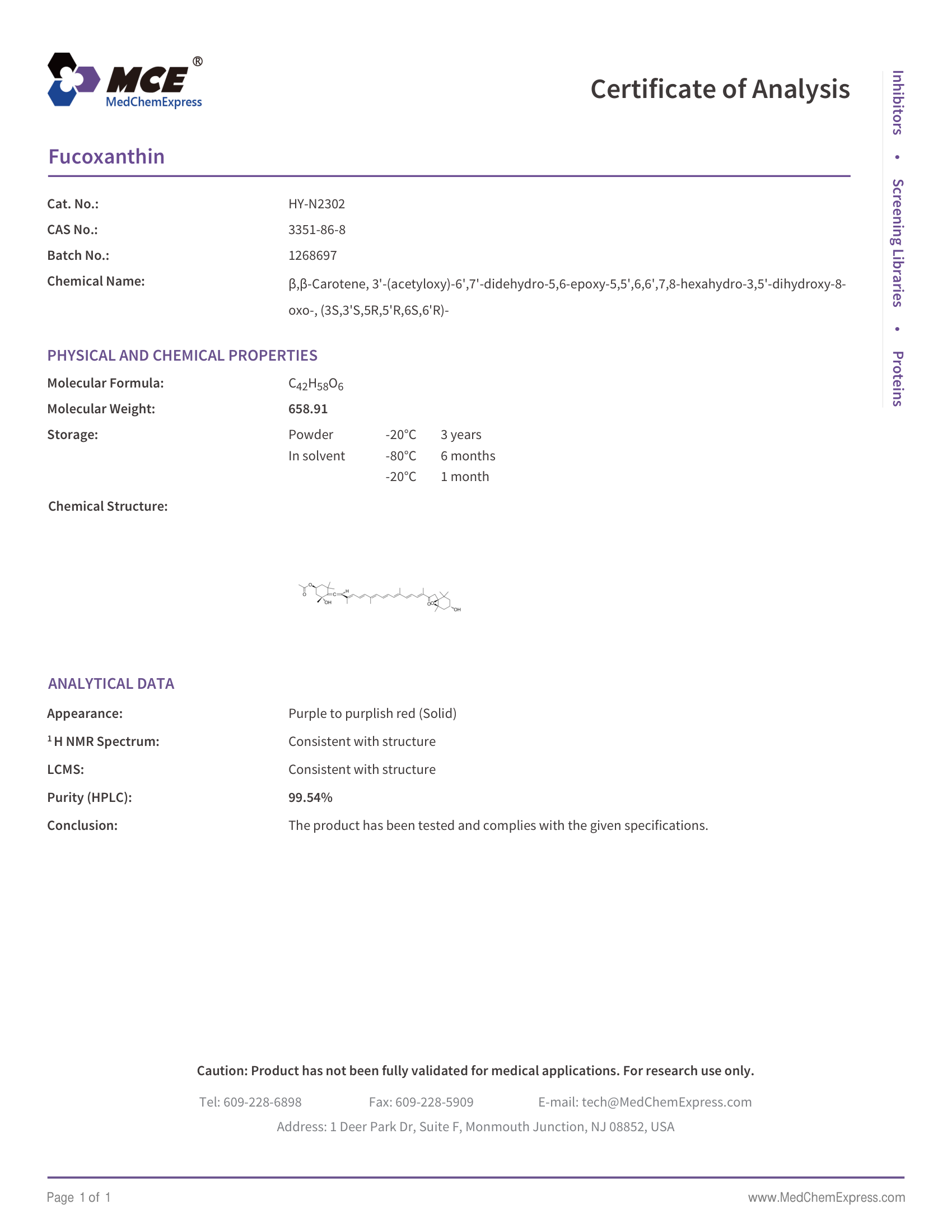

Figure S1. Certificate of analysis for fucoxanthin.

Manufacturer's certificate of analysis for the fucoxanthin used in this study (cat. no. HY-N2302, batch no. 1268697, MedChemExpress), reporting a purity of 99.54% by HPLC, with ¹H NMR and LCMS consistent with the assigned structure. A single batch was used for all experiments reported in this study.

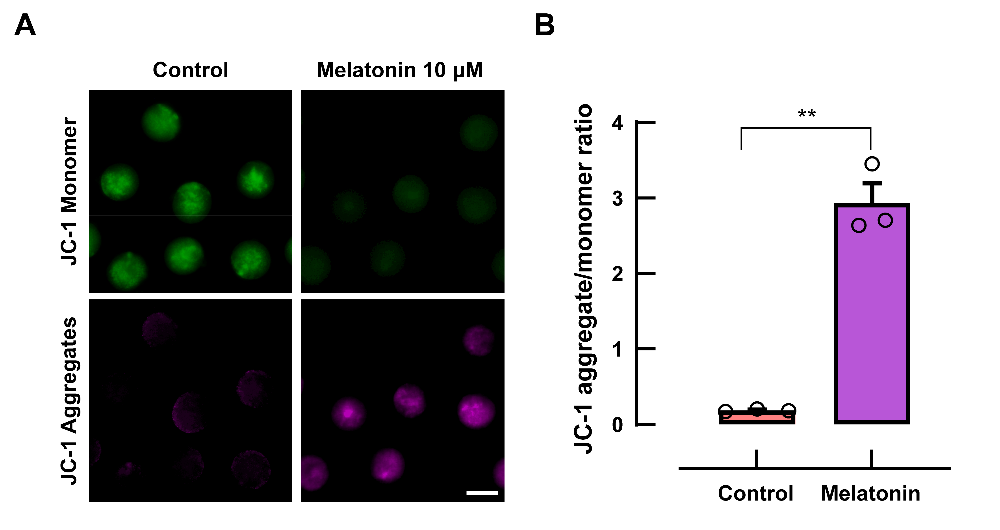

Figure S2. Melatonin restores mitochondrial membrane potential during postovulatory aging, validating the JC-1 assay as a positive-control readout.

(A) Representative fluorescence images of JC-1 staining after 12 h of POA. Green indicates JC-1 monomers (depolarized mitochondria) and magenta indicates JC-1 aggregates (polarized mitochondria). Melatonin (10 µM) was added to intact cumulus–oocyte complexes at the onset of POA culture, and Control received vehicle only. The final DMSO concentration was adjusted to 0.1% (v/v) in both groups. Oocytes were denuded immediately before imaging. Scale bar, 100 µm.

(B) Quantification of mitochondrial membrane potential (ΔΨm) from the images shown in (A), expressed as the ratio of JC-1 aggregate to monomer fluorescence and quantified as described in Section 2.7 of the main text. Data are mean ± SEM of three independent replicates (n = 3), with 6–23 oocytes analyzed per group in each replicate, and each open circle represents one independent replicate. Melatonin, an antioxidant with established protective activity in this model, was included in the same experiments as those shown in Fig. 1 to confirm that the assay could detect an improvement in ΔΨm; the Control group is therefore shared with Fig. 1D. Paired *t*-test with replicates matched by batch (***p* < 0.01).

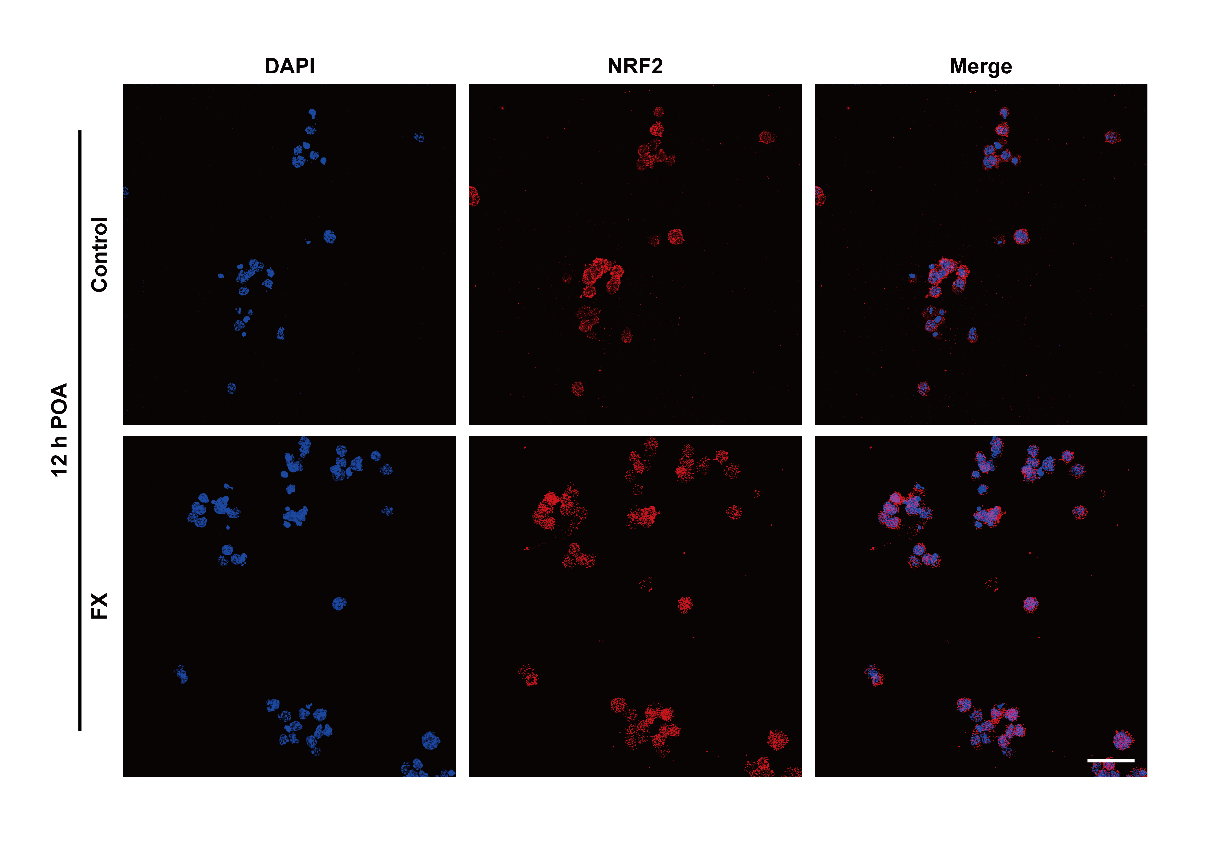

**Figure S3. Low-magnification fields of NRF2 immunofluorescence in cumulus cells.** Representative fields from which the single cells shown in Fig. 4A were cropped. Cumulus cells were dispersed from cumulus–oocyte complexes after 12 h of POA, attached to adhesive slides, and immunostained for NRF2 (red) with DAPI counterstaining (blue). Top, Control; bottom, FX 5 µM. Quantification was performed on all cells within such fields, as described in Section 2.7 of the main text. Scale bar, 25 µm.

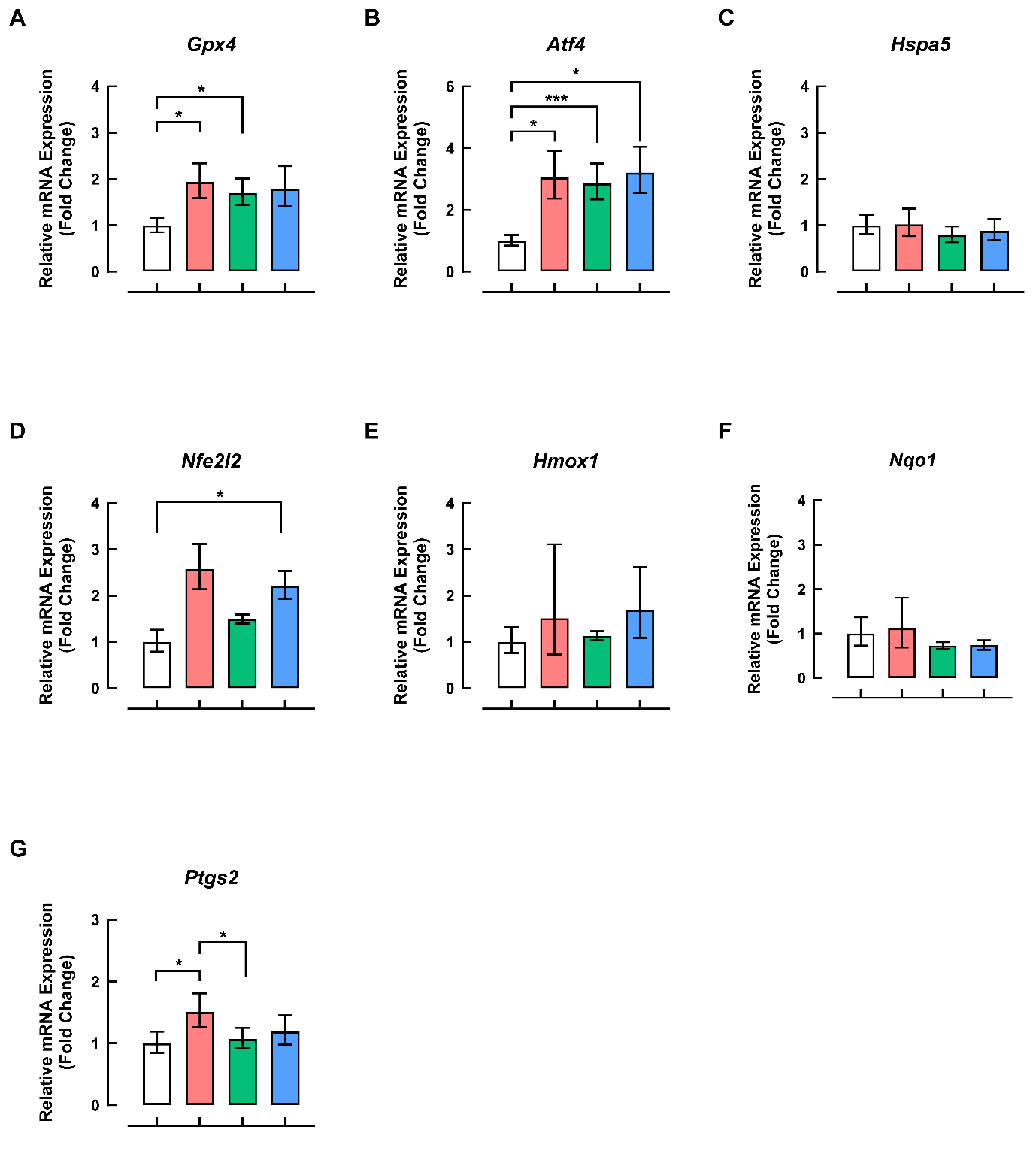

**Figure S4. Postovulatory aging activates a transcriptional stress response in cumulus cells that is not further modified by fucoxanthin.**

Relative mRNA expression in cumulus cells isolated after 12 h of POA, measured by quantitative PCR. (A) *Gpx4*, (B) *Atf4* and (C) *Hspa5*, representing the antioxidant and integrated stress responses. (D) *Nfe2l2*, (E) *Hmox1* and (F) *Nqo1*, representing the NRF2 axis and its two canonical target genes. (G) *Ptgs2*, encoding cyclooxygenase-2. In every panel the bars are, from left to right, Fresh (white), Control (red), FX (green) and AX (blue).

Cumulus–oocyte complexes were aged for 12 h in M16 medium containing vehicle only (Control), fucoxanthin (FX, 5 µM) or astaxanthin (AX, 5 µM), with the final DMSO concentration adjusted to 0.1% (v/v) in all groups. Complexes collected at 14 h post-hCG and processed without aging culture served as the non-aged reference group (Fresh). Cumulus cells were dissociated at the end of the aging period and pooled within each replicate. Expression was calculated by the 2^−ΔΔCt^ method with *H2az1* (formerly *H2afz*) as the internal control and normalized to the Fresh group. Data are mean ± SEM of three independent replicates (n = 3). Repeated-measures one-way ANOVA with Tukey's multiple comparisons test (**p* < 0.05; ******p <* 0.001); comparisons without an indicated bracket were not statistically significant. Primer sequences are listed in Table S1.

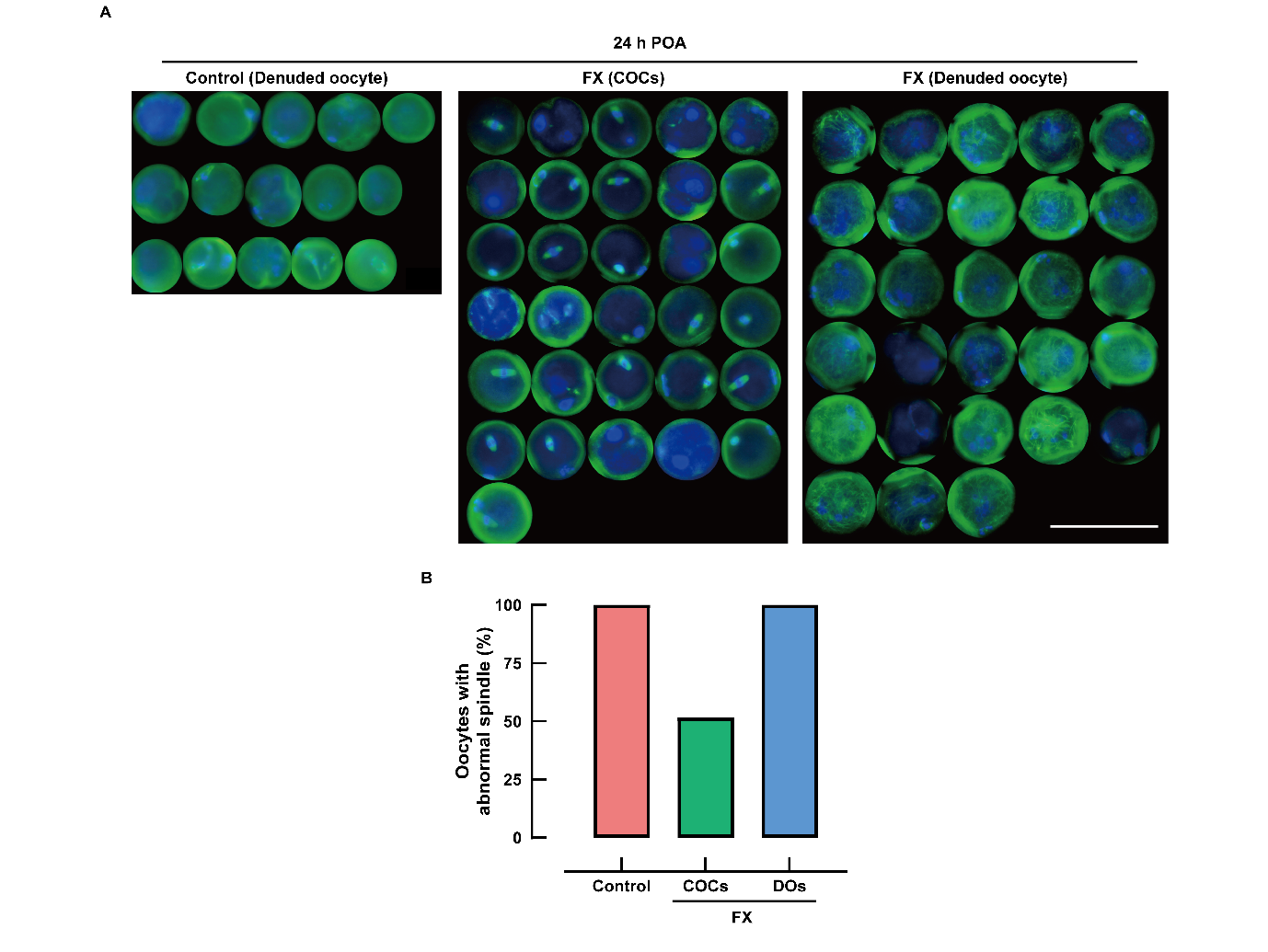

**Figure S5. Spindle organization in fucoxanthin-treated oocytes aged within intact cumulus–oocyte complexes or after denudation (preliminary experiment).**

(A) Meiotic spindle morphology after 24 h of POA: α-tubulin (green), DAPI (blue). Intact cumulus–oocyte complexes (COCs) or oocytes denuded before culture (DOs) were aged in M16 medium containing vehicle only (Control) or fucoxanthin (FX, 5 µM), with the final DMSO concentration adjusted to 0.1% (v/v) in all groups. Oocytes in the COC group were denuded immediately before fixation. Images of individual oocytes were cropped from multiple fields of the same experiment and assembled for presentation. Scale bar, 100 µm. (B) Proportion of oocytes with abnormal spindles, scored by an observer blinded to treatment using the criteria defined in Section 2.7 of the main text. All oocytes were abnormal in the Control group and in the FX (DOs) group, whereas about half were abnormal in the FX (COCs) group. Note that the control arm of this experiment comprised denuded oocytes only, so the comparison that isolates the contribution of the cumulus investment is FX (COCs) versus FX (DOs). Data are from a single experiment (n = 1; 15, 31 and 28 oocytes for Control, FX (COCs) and FX (DOs), respectively) and are therefore presented without statistical analysis.
